# The ComRS-SigX pathway regulates natural transformation in *Streptococcus ferus*

**DOI:** 10.1101/2023.03.06.531454

**Authors:** Britta E. Rued, Michael J. Federle

**Affiliations:** Department of Pharmaceutical Sciences. University of Illinois at Chicago, Chicago, Illinois, USA

## Abstract

The ability to take up and incorporate foreign DNA via natural transformation is a well-known characteristic of some species of *Streptococcus,* and is a mechanism that rapidly allows for the acquisition of antibacterial resistance. Here, we describe that the understudied species *Streptococcus ferus* is also capable of natural transformation and uses a system analogous to that identified in *Streptococcus mutans*. *S. mutans* natural transformation is under the control of the alternative sigma factor *sigX* (also known as *comX*), whose expression is induced by two types of peptide signals: CSP (<u>c</u>ompetence <u>s</u>timulating <u>p</u>eptide, encoded by *comC*) and XIP (*sig<u>X</u>*-inducing <u>p</u>eptide, encoded by *comS*). These systems induce competence via either the two-component signal-transduction system ComDE or the RRNPP transcriptional regulator ComR, respectively. Protein and nucleotide homology searches identified putative orthologs of *comRS* and *sigX* in *S. ferus*, but not homologs of *S. mutans blpRH* (also known as *comDE*). We demonstrate that natural transformation in *S. ferus* is induced by a small, double-tryptophan containing competence-inducing peptide (XIP), akin to that of *S. mutans*, and requires the presence of the *comR* and *sigX* orthologs for efficient transformation. Additionally, we find that natural transformation is induced in *S. ferus* by both the native XIP and the XIP variant of *S. mutans*, implying that crosstalk between the two species is possible. This process has been harnessed to construct gene deletions in *S. ferus* and provides a method to genetically manipulate this understudied species.

**IMPORTANCE:** Natural transformation is the process by which bacteria take up DNA and allows for acquisition of new genetic traits, including those involved in antibiotic resistance. This study demonstrates that the understudied species *Streptococcus ferus* is capable of natural transformation using a peptide-pheromone system like that previously identified in *Streptococcus mutans* and provides a framework for future studies concerning this organism.

## INTRODUCTION

Natural transformation is the process by which bacteria take up and incorporate DNA from their environment (1–3). It can lead to drastic changes in genetic content of an organism and can result in acquiring advantageous traits. Of particular importance in clinical situations, rapid evolution through this form of horizontal gene transfer can enhance an organism’s virulence or resistance to antimicrobial agents (2–4). The bacterial physiological state allowing for natural transformation, known as competence, is stimulated in *Streptococcus pneumoniae* by the secreted pheromone *<u>c</u>*ompetence *<u>s</u>*timulating *<u>p</u>*eptide (CSP; encoded by *comC*), which is sensed by a membrane sensor kinase/response regulator pair called ComDE (5, 6). This results in induction of early competence genes, one being *sigX* (otherwise known as *comX*), the alternative sigma factor required for RNA polymerase to express late competence genes (7–10). In many streptococcal species, induction of this system leads to expression of genes involved in fratricidal behavior or bacteriocin production (1–3, 11). This process is thought to contribute to bacterial competition and allow for closely related species to “steal” DNA from other species they might encounter in their native niche. Of course, this process can also be exploited in the laboratory to enhance genetic manipulation for deeper studies.

Interestingly, *S. mutans* and related species, possess not just a ComCDE-like system (recently redesignated BlpRH for its involvement in bacteriocin production and conservation of genomic architecture to other BlpRH systems) (11–13), but also a separate regulatory entity termed ComRS of the Rgg/Rap/Npr/Plc/Prg (RRNPP) family that directly induces competence. This pathway relies on the production and uptake of the signaling pheromone termed XIP (*<u>si</u>*gX *i*nducible *<u>p</u>*eptide) (14, 15). Upon uptake of XIP via the Opp (Ami) transporter, XIP binds to ComR and directly induces transcription of *comS* (encoding the XIP pre-peptide) and *sigX* (16–18). As in *S. pneumoniae*, SigX assembles with RNA polymerase to transcribe downstream competence genes to induce natural transformation (7, 8). The ComRS system was discovered relatively recently compared to the more longstanding research of the ComCDE and BlpRH systems. In this short period of time, multiple groups have investigated factors such as cross-talk between ComCDE/BlpRH, ComR, and other regulators, intracellular signaling, the structure of ComR, and the effects of XIP induction on *S. mutans* physiology, as reviewed in (19, 20). This work led to additional understanding about the various ways competence can be induced or modulated in *S. mutans*.

While related to *S. mutans*, *S. ferus* is a relatively understudied organism. It was originally described as a “Mutans-like” streptococcus isolated from the oral cavities of wild rats (21) and later found in oral cavities of pigs (22, 23). Despite being similar to *S. mutans* in its ability to colonize oral cavities, it does not cause caries in rodent models, although it can form dental biofilms (plaque) and has acidogenic potential (21). Recently, we described two ribosomally encoded and post-translationally modified peptides (RiPPs), called Triglysins, that are produced by *S. mutans* and *S. ferus*, contain antimicrobial activities, and are regulated by another RRNPP quorum-sensing system (24). While our studies benefited from genetic manipulation of *S. mutans* made possible through the harnessing of natural transformation, methodologies to engineer genetic variants in *S. ferus* were not described in the literature. Here, we characterize a ComRS signaling pathway in *S. ferus* and demonstrate robust induction of natural transformation and genetic modification.

## RESULTS

### Identification of a ComRS system in *S. ferus*

To examine if *S. ferus*, like other streptococci, had the potential for natural competence we first searched for the presence of known competence system orthologs. Two well-characterized systems that induce competence in *S. mutans* are the ComRS and BlpRH (formerly known as ComCDE) systems. *S. ferus* is related to *S. mutans*, although it is distinct in certain phenotypes such as caries potential and acid resistance (21, 22). We first performed a protein BLAST search (25) using the *S. mutans* ComR, its homolog SMU_381c, *S. mutans* PdrA (another known Rgg transcriptional regulator) and BlpRH to identify if orthologs also existed in *S. ferus*. No orthologs of the *S. mutans* BlpRH system were found; however, we identified two proteins with similarity to ComR and SMU_381c in *S. ferus* (Table 1). The two orthologs corresponded to genes labeled *A3GY_RS0108865* and *A3GY_RS0106270*, with RS0108865 having 61% identity to *S. mutans* ComR and RS0106270 having 44% identity to *S. mutans* SMU_381c, a previously identified ComR homolog (14). Due to the higher percent identity and additional factors discussed below, we designated A3GY_RS0108865 as *comR* and A3GY_RS0106270 as *comR2*. The amino acid alignment between these and other identified ComR/Rgg orthologs in *S. ferus* is shown in Fig. S1A. Two other Rgg orthologs were identified by BLAST analysis of the *S. mutans* PdrA sequence: A3GY_RS0105975, which we designated *rggA*, and *S. ferus* PdrA (A3GY_RS0100490), an Rgg ortholog that has been previously identified and regulates the production of the RaS-RiPP tryglysin A (24). These two Rgg orthologs had lower percent identity to *S. mutans* ComR, 17% and 18% respectively (Table 1). Overall, *S. ferus* possesses 4 ComR/Rgg-like proteins.

**TABLE 1.** Amino acid identity between ComR/Rgg orthologs identified in S. ferus and S. mutans.

| Locus in *S. mutans* or *S. ferus* DSM20646<sup>a</sup> | Gene name | *S. mutans* ComR (SMU_61) | *S. mutans* ComR2 (SMU_381c) | *S. mutans* PdrA (SMU_1509) |
| --- | --- | --- | --- | --- |
| SMU_61 | *comR* | N/A<sup>b</sup> | 32% | 20% |
| SMU_381c | *comR2* | 32% | N/A<sup>b</sup> | 17% |
| SMU_1509 | *pdrA* | 20% | 17% | N/A<sup>b</sup> |
| A3GY_RS0108865<sup>c</sup> | *comR* | 61% | 32% | 18% |
| A3GY_RS0106270<sup>c</sup> | *comR2* | 34% | 44% | 17% |
| A3GY_RS0105975<sup>c</sup> | *rggA* | 17% | 14% | 27% |
| A3GY_RS0100490<sup>c</sup> | *pdrA* | 18% | 15% | 65% |

<sup>a</sup>Amino acid sequence for each protein was obtained from *S. mutans* UA159 or *S. ferus* DSM20646. See *Materials and Methods* for further details.
<sup>b</sup>N/A: Not applicable.
<sup>c</sup>Annotated as the following loci in NCTC12278 assembly: *comR*, DQL21\_RS00510; *comR2*, DQL21\_RS08780; *rggA*, DQL21\_RS09080; *pdrA*, DQL21\_RS02900.

We then examined the genetic architecture around the putative ComR/Rgg orthologs, reasoning that the putative *comS* open reading frame would be encoded nearby the identified *comR*, as in other comRS systems. We identified a small open reading frame encoding 13 amino acids that was 76 bp downstream of the stop codon of *comR* (Fig. 1A). This small open reading frame (sORF) was predicted to encode for the short peptide MFRFLTGLSWWGL and possesses the characteristic double-tryptophan motif previously identified in *S. mutans* XIP and XIPs from other ComRS systems (14) (Fig. 1D, Table 2). We compared the predicted sORF to the identified *comS* from *S. mutans* and found that the final 7 amino acids were highly conserved, with the only differences being an Asp to Ser and Ser to Gly at the 9^th^ and 12^th^ amino acids in this sequence (Fig. 1D, Table 2). As such, we designated the small open reading frame as the putative *S. ferus comS*. Examining the region surrounding A3GY_RS0106270 (*comR2*) revealed five predicted sORFs, but only two aligned with a promoter sequence like the identified *sigX* consensus sequence in *S. mutans* (Fig. 1A, 1C, Table 2) (14). We gave the two small ORFs following *comR2* a general designation of *orf01* and *orf02* and noted that both appeared to be enriched for hydrophobic or branched chain amino acids, like other ComR/Rgg type signaling peptides.

**FIGURE 1:**
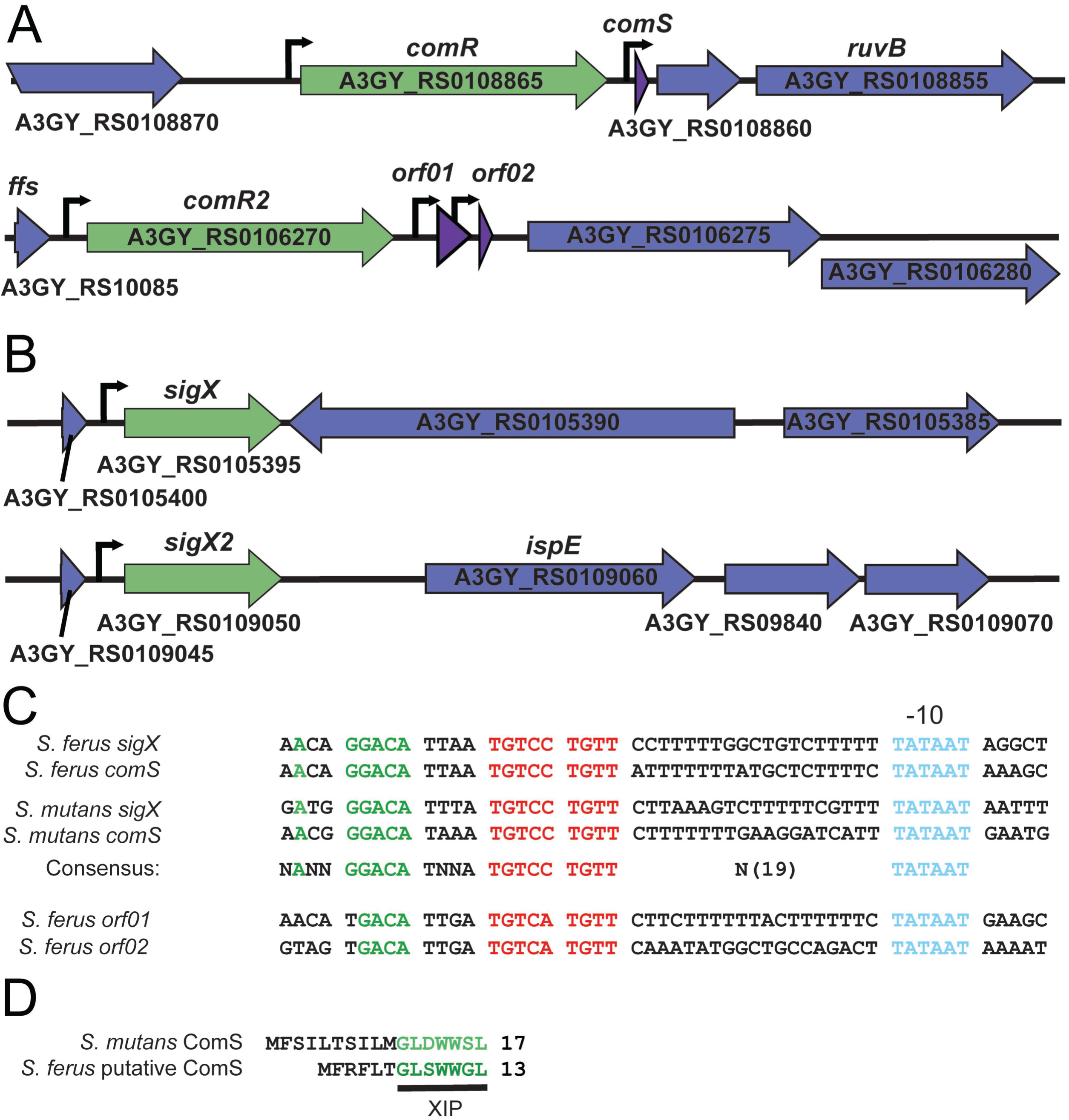
Identification of a ComRS system and related ComR ortholog in *S. ferus*. A) Locus arrangement of *comRS* and *comR2* in *S. ferus* DSM20646. B) Locus arrangement of *sigX* and *sigX2* in *S. ferus* DSM20646. C) Examination of the sequences upstream of *S. ferus sigX*s (*comX/2,* A3GY_RS0105395 and A3GY_RS0109050), *comS*, *orf01* and *orf02* versus *S. mutans* UA159 *sigX* (SMU_61) and *comS*. Highlighted bases: blue, conserved −10 hexamer; green and red, other conserved bases. D) Amino acid alignment of the *S. mutans* ComS and the *S. ferus* putative ComS. Green underlined letters indicate the predicted XIP sequence.

**TABLE 2.**
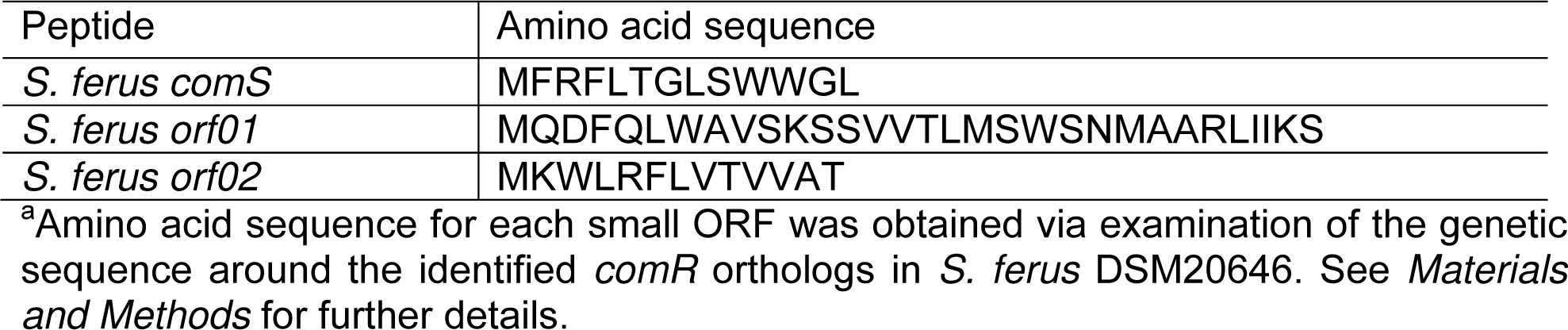
Amino acid sequence of putative small ORFs identified in S. ferus.^a^

**TABLE 3.** Amino acid sequence of peptides used in this publication.^a^

| Peptide | Amino acid sequence |
| --- | --- |
| *S. ferus* XIP | GLDWWSL |
| *S. mutans* XIP | GLSWWGL |
| *S. mutans* reverse SHP | GGGIIIITE |

<sup>a</sup>XIP *S. mutans* was synthesized by NeoScientific, whereas XIP *S. ferus* and reverse SHP *S. mutans* were synthesized by ABClonal. See *Materials and Methods* for further details.

We then performed a BLASTp search using the amino acid sequence for *S. mutans* SigX, reasoning that this would help us identify if an intact ComRS system existed in *S. ferus*, as SigX functions as the alternative sigma factor required for induction of competence in most streptococci (3, 7). Streptococci have typically been found to have one or two copies of *sigX* (7, 14, 26). In species such as *Streptococcus pneumoniae* with two *sigX* genes, the presence of either copy is sufficient to allow transformation at wild-type levels (7). Within the two available *S. ferus* genomes on NCBI, A3GY_RS0105395 and A3GY_RS0109050 (which are listed as replicate strains according to the U.K. Health Security Agency (https://www.culturecollections.org.uk/)), we found two *sigX* paralogs, differing only at amino acid 103 as a Gly or Cys residue; we designate these *sigX* and *sigX2*, respectfully (Fig. 1B, S1B). The genes have similar 5’ flanking regions, with an apparent duplication of a tRNA-Asn, and a short 3’ flanking region consisting of 39 bp, after which the sequences diverge. This genetic architecture around the identified *sigX* orthologs mimics that reported for other streptococci (14).

Upon identifying the two SigX orthologs, we next examined the putative promoter regions of *S. ferus comS*, *sigX*, *sigX2*, *orf01*, and *orf02* to determine if a promoter structure like that of *S. mutans* or *S. salivarius sigX* P1 or P2 promoter was present (14). This and related sequences have been identified to be required for ComR control of *comS* and *sigX* expression in *S. mutans*, *S. salivarius*, and other streptococci (14, 15, 27). Directly upstream of the putative *comS*, *sigX*, and *sigX2*, we identified a motif almost completely identical to that of the *S. mutans* P1 promoter (Fig. 1C). This motif conserves the inverted repeat previously identified in this promoter as well as the positioning of the repeated 19 nucleotides upstream of the canonical −10 hexamer (14). Interestingly, for *orf01* and *orf02* the promoter pattern was also highly similar; with the exception of the first nucleotide of the inverted repeat, the rest of the core sequence and spacing were conserved (Fig. 1C). This suggests that if expressed, these sORFs are likely to be controlled by ComR2 in a similar manner as ComR controls *comS*. We did not pursue this circuit for further examination, reasoning that the primary *comRS* identified was more likely involved in competence induction in *S. ferus*, due to previous findings in *S. mutans* (14). Instead, we focused on characterizing the putative ComRS circuit to harness this for genetic manipulation of *S. ferus*.

### Induction of natural transformation in *S. ferus*

We first examined if *S. ferus* could be rendered transformable by the addition of synthetic *S. ferus* or *S. mutans* XIP. We reasoned that since the two XIPs were so similar in sequence at the C-terminus, they would likely both induce competence in *S. ferus*. *S. ferus* cultures grown in a peptide-free, chemically defined medium (CDM) and supplied with either a linear DNA amplicon (Δ*comR*::*spec*) or a closed-circular plasmid (pLZ12-*spec*), were treated with XIP. Transformants were obtained after stimulation with either version of XIP, with frequencies of 0.1 to 2% seen with *S. ferus* XIP, and 0.04-0.4% for the *S. mutans* version (Fig. S2A-B). Efficiencies were significantly lower for plasmid vs linear DNA, an observation that has previously been made for other bacterial species (Fig. S2A) (28, 29). From these experiments, we find that both *S. ferus* and *S. mutans* XIP orthologs are capable of inducing transformation in *S. ferus*, indicating that crosstalk between these two species might occur.

As we occasionally observed transformants in wildtype *S. ferus* in the absence of exogenously supplied XIP, although infrequent and highly variable (Fig. 2B, third column), we reasoned that wild type *S. ferus* produced XIP under the defined growth conditions. To investigate the contribution that endogenously produced XIP would have on competence, we generated a strain [*comS*(GGG)] for which the start codon of *comS* was mutated (ATG → GGG) and unable to support translation initiation. We assessed transformation efficiency when synthetic XIP was provided and found that frequencies were slightly lower than seen in wildtype responses to *S. ferus* XIP, but the decrease was not significant (Fig. 2B, Fig. S2C). No transformants were observed in the *comS*(GGG) strain without supplemented XIP (Fig. 2B). We then tested the ability of spent culture supernatants from early stationary phase cultures of wildtype to induce competence in the *comS*(GGG) strain but no transformants were observed (Fig. S2F). Therefore, we conclude that endogenous production of XIP by *S. ferus* is limited, but likely to occur in lab conditions; however, capturing XIP from supernatants for purposes of stimulating heterologous cultures will likely require a means to concentrate the signal.

**FIGURE 2:**
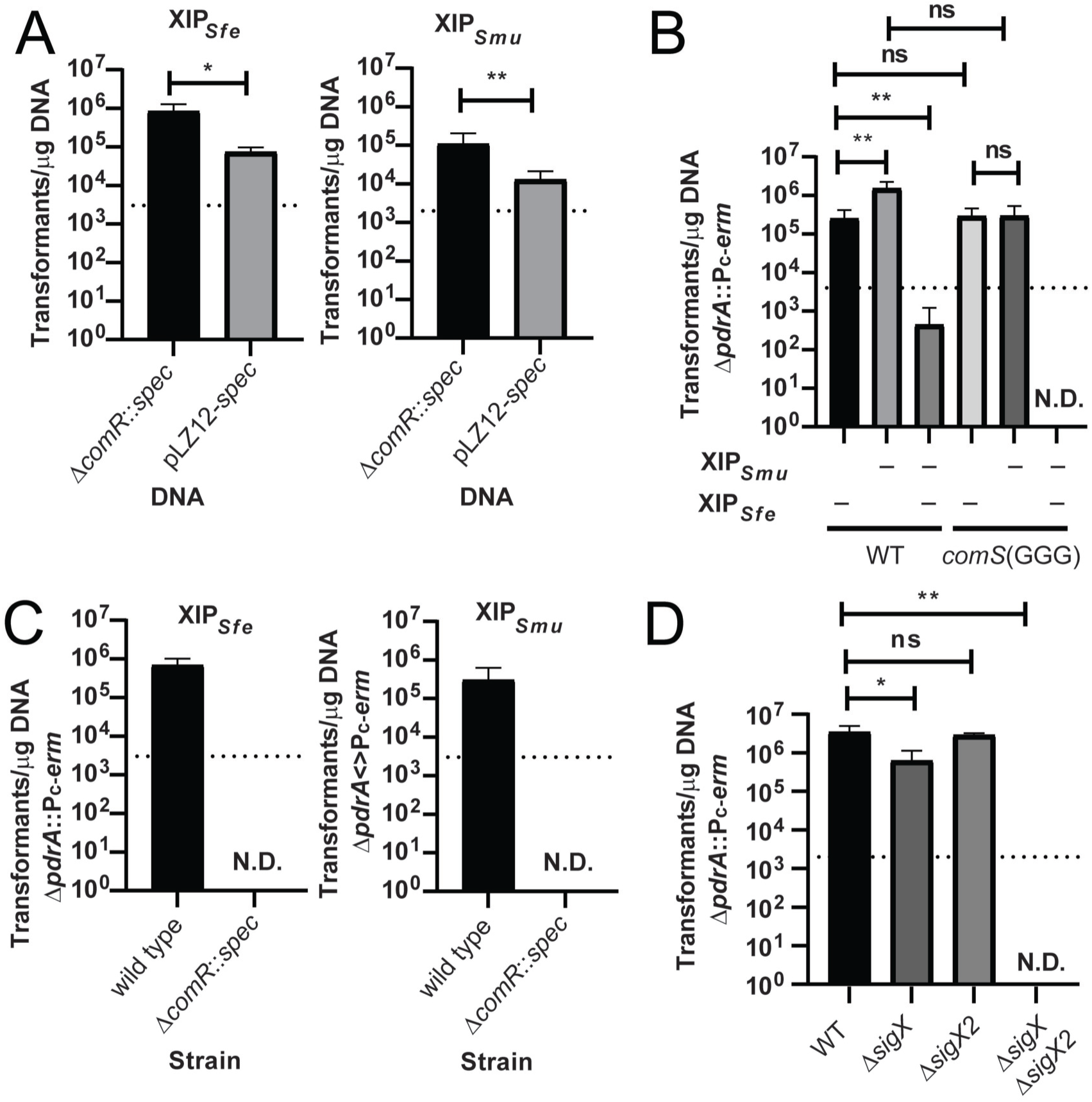
Characterization of *S. ferus* transformation in wild-type *S. ferus* and competence gene mutants. A) Transformation efficiency of *S. ferus* with a linear amplicon (Δ*comR*::*spec)* or plasmid (pLZ12-*spec*) with 1 µM *S. ferus* or *S. mutans* XIP, as indicated above the graph. DNA used is indicated below the graph, and the dotted line indicates the limit of detection. This graph is a summary of three independent experiments. B) Transformation efficiency of *S. ferus* wild-type and *comS*(GGG) strains with 1 µM *S. ferus* or *S. mutans* XIP, as indicated below the graph. 50 ng of linear Δ*pdrA*::P_c_-*erm* were used for each transformation. N.D. indicates not detected, as no detectable colonies were observed on transformation plates. This experiment was performed three times with similar results. C) Transformation efficiency of *S. ferus* wild-type vs. Δ*comR* with 1 µM *S. ferus* or *S. mutans* XIP, as indicated above the graph. Strain used is indicated below the graph, and the dotted line indicates the limit of detection. 100 ng of linear Δ*pdrA*::P_c_-*erm* were used for each transformation. N.D. indicates not detected, as no detectable colonies were observed on transformation plates. This graph is a summary of three independent experiments. D) Transformation efficiency of *S. ferus* Δ*sigX*, Δ*sigX2*, Δ*sigX* Δ*sigX2* vs. wild-type *S. ferus* with 50 ng of a linear amplicon (Δ*pdrA::*P*_c_*-*erm*) and 1 µM *S. ferus* XIP. The dotted line on the graph indicates limit of detection. N.D. indicates not detected, as no detectable colonies were observed on transformation plates. This experiment was performed three times with similar results.

We next interrogated the role of ComR and SigX for competence induction, as it is essential for other species (15, 27). We deleted the *comR* gene using an in-frame insertion of a spectinomycin cassette in *comR*. In the absence of *comR*, transformants were not detected (Fig. 2C, S2D). The necessity of the two *sigX* genes was determined by constructing deletions of *sigX*, *sigX2*, and a combined *sigX sigX2* deletion, and examining transformation in these backgrounds. With a linear amplicon, we observed that neither deletion of *sigX* nor *sigX2* alone prevented *S. ferus* from undergoing transformation (Fig. 2D), although the *sigX* strain did show significantly reduced transformation frequency (Fig. S2E). Like findings reported from *S. pneumoniae* (7), we found that only the deletion of both copies of *sigX* resulted in abrogation of transformation (Fig. 2D, Fig. S2E).

### Impact of time, XIP and DNA concentration on transformation in *S. ferus*

To further characterize the limits of natural transformation in *S. ferus*, we first examined the impact of varying the concentration of XIP on transformation efficiency. For a linear DNA construct with approximately 1 kB flanking regions of homologous sequence, we observed maximal transformation efficiency occurring between 100 nM and 1 µM *S. ferus* XIP (Fig. 3A). To determine how DNA concentration impacted the number of transformants arising, XIP was kept constant at 1 µM and the concentration of linear DNA was varied. No transformation occurred without the addition of DNA and a minimum of 5 ng DNA was required to observe reproducible transformation events, with saturation being seen starting at 100 ng (Fig. 3B). Efficiency (transformants per ug DNA) remained consistent at all tested concentrations. Finally, we evaluated the amount of time required after XIP treatment to recover transformants. Mid-exponential growth phase cultures were stimulated with 1 µM XIP and provided with 50 ng of DNA, after which samples were plated on selective agar periodically. By 15 minutes after XIP stimulation, the recovery of resistant colonies was emergent, and by 60 minutes the peak rate was reached and held steady for at least 3 hours (Fig. 3C).

**FIGURE 3:**
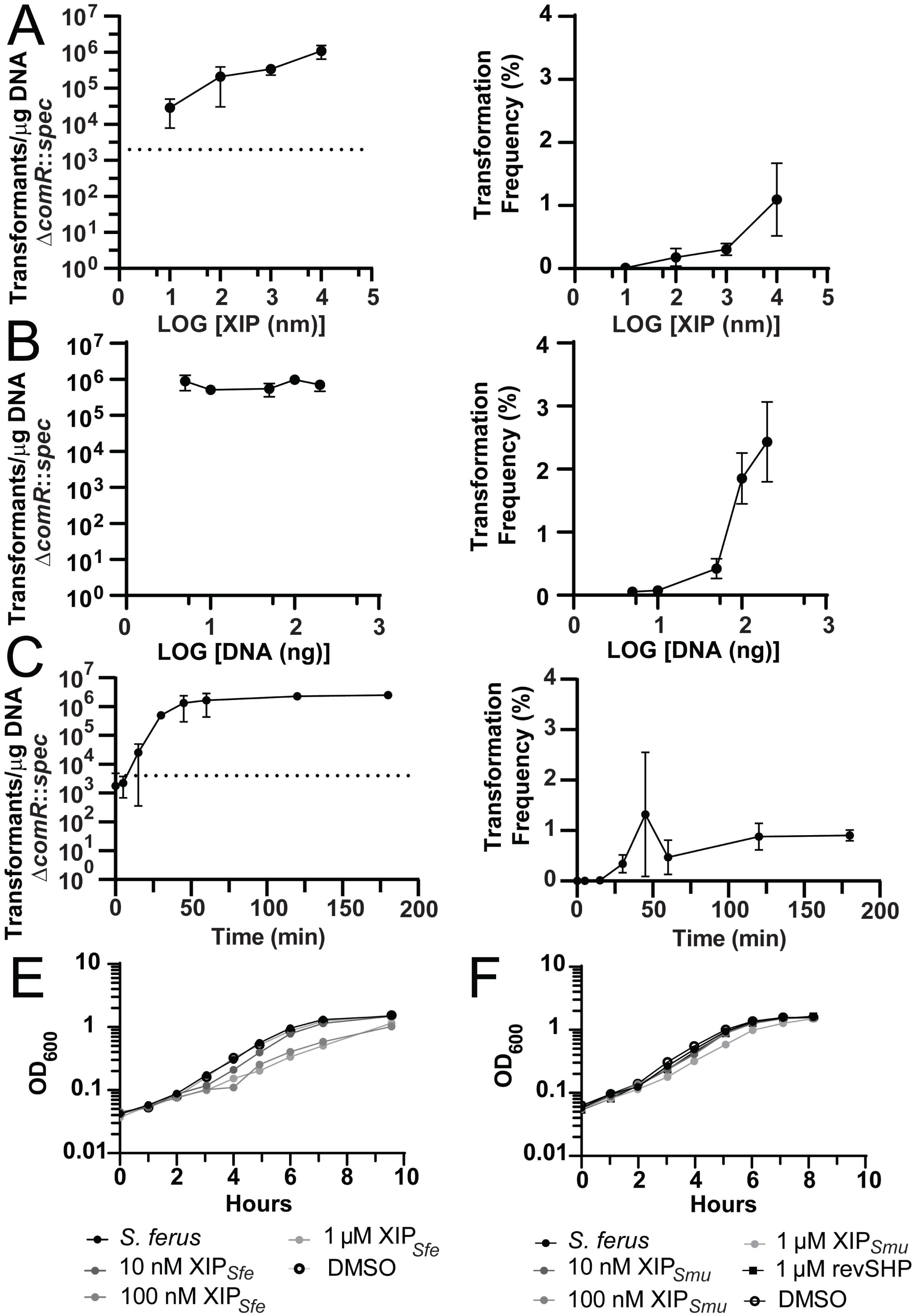
Examination of the effect of XIP concentration, DNA concentration, and time on *S. ferus* transformation, as well as XIPs on *S. ferus* growth. For further experimental details, see *Materials and Methods*. A) Graphs of transformation efficiency and transformation frequency vs. *S. ferus* XIP concentration for 100 ng of a linear DNA amplicon (Δ*comR*::*spec*). This experiment was performed three times with similar results. B) Graphs of transformation efficiency and transformation frequency vs. DNA concentration of a linear DNA amplicon (Δ*comR::spec*) with 1 µM *S. ferus* XIP. This experiment was performed three times with similar results. C) Graphs of transformation efficiency and frequency over time for 50 ng of a linear DNA amplicon (Δ*comR*::*spec*). This experiment was performed twice with similar results. E) Growth curve of *S. ferus* in CDM with increasing amounts of *S. ferus* XIP or equivalent DMSO carrier vehicle. Strains and conditions are indicated in the legend below the graph. This experiment was performed three times with similar results. F) Growth curve of *S. ferus* in CDM with increasing amounts of *S. mutans* XIP or equivalent DMSO carrier vehicle. This experiment was performed three times with similar results.

### Impact of XIP and competence on growth rate

The impact that addition of exogenous XIP had on *S. ferus* growth rates was determined using titrations of synthetic *S. ferus* and *S. mutans* XIP. 100 nM and 1 µM *S. ferus* XIP significantly increased the doubling time of wild-type bacteria (Fig. 3E, Table S2), whereas *S. mutans* XIP did not have a statistically significant effect, although a slight reduction in the growth rate was observed at the highest concentration of peptide (Fig. 3F, Table S2). This result correlates with the higher specific activity of the *S. ferus* XIP. Controls with the vehicle DMSO or equivalent concentrations of a reverse SHP peptide from *S. mutans* known to be ineffective for signaling (24) did not alter the growth rate. Deletion of *comR* resulted in a strain that slowed doubling after 4 hours of growth and this defect was not exacerbated by the addition of XIP (Fig. S3A, Table S2). Disruption of *sigX* and *sigX2* had no significant effect on growth, but interestingly, the XIP dependent increase in doubling time was no longer significant when *sigX* was disrupted (Fig. S3B, Table S2 second column). The double deletion of *sigX* and *sigX2* also resulted in a strain that did not have a statistically significant increase in doubling time with XIP (Table S2, Fig. S3C).

### Response of *comR*, *comS*, and *sigX* to XIP

Finally, we measured the transcriptional response of *comR, comS*, and *sigX* to the addition of XIP by creating transcriptional fusions of the promoter regions of each gene to *luxAB*. We observed that P*_comR_* transcription was expressed constantly regardless of the presence of XIP in both CDM and THY (peptide-rich) media (Fig. 4A, 4E). A strong response to XIP was seen for P*_sigX_* in CDM; however the reporter was also induced in the absence of XIP, albeit at higher cell densities, indicating that natural induction of competence was occurring through endogenous production of XIP (Fig. 4B). This is consistent with the expression profiles of the P*_comS_* reporters, where rapid stimulation was seen upon addition of both XIP versions, but induction was also seen without exogenous XIP stimulation (Fig. 4C-D). Interestingly, autoinduction was not seen for either P*_sigX_* or P*_comS_* when cultures were grown in THY (Fig. 4E-F).

**FIGURE 4:**
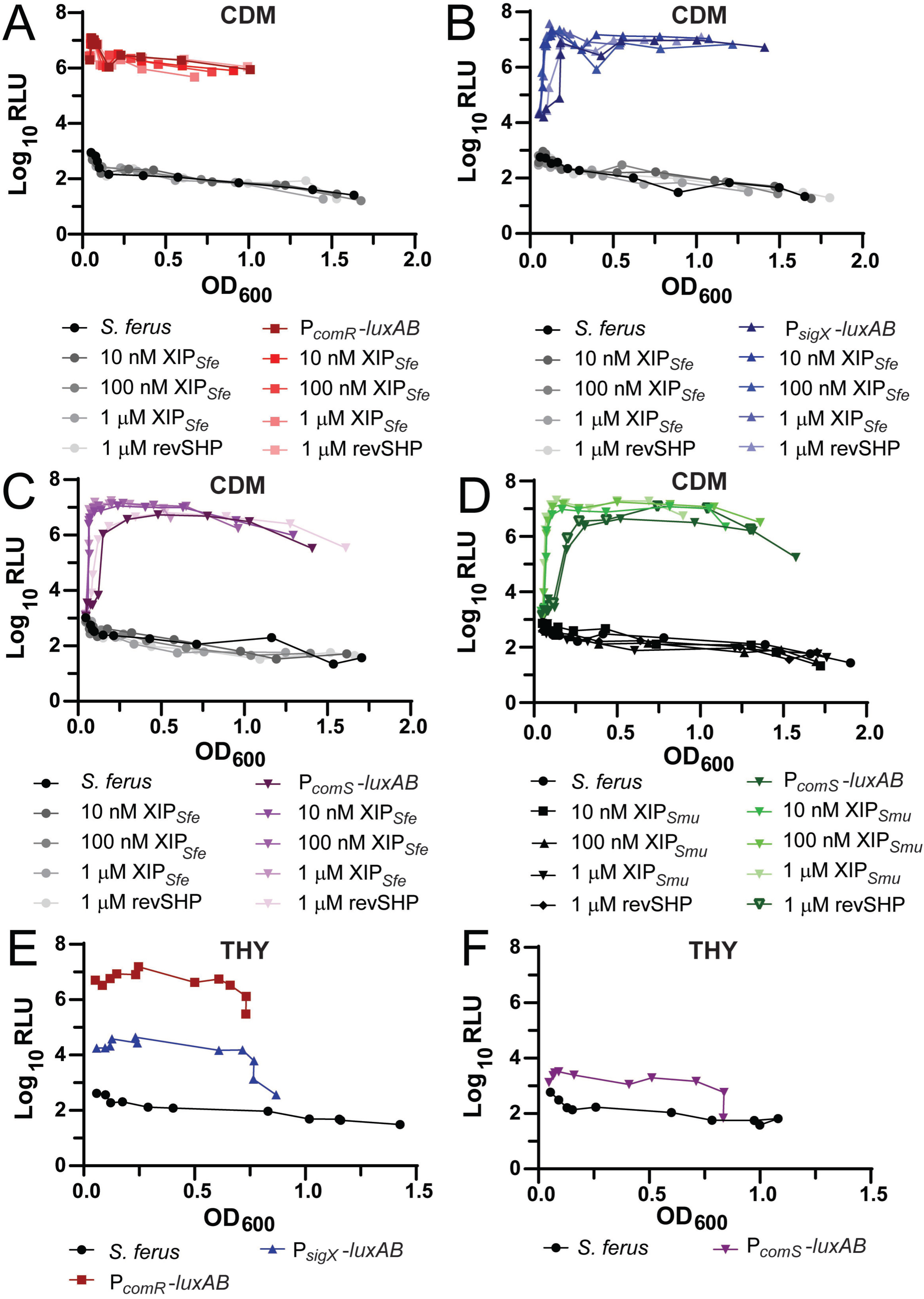
Response of *comR*, *comS*, and *sigX* luciferase reporters to XIP addition in chemically defined media (CDM) or THY, as indicated above each graph. Conditions and strains are indicated in the legend below each graph, and each experiment was performed three times with similar results. XIP or revSHP concentrations added to cultures were approximate. A) P*_comR_*-*luxAB* reporter response to increasing *S. ferus* XIP. B) P*_sigX_*-*luxAB* reporter response to increasing *S. ferus* XIP. C) P*_comS_*-*luxAB* reporter response to increasing *S. ferus* XIP. D) P*_comS_*-*luxAB* reporter response to increasing *S. mutans* XIP. E) P*_comR_*-*luxAB* and P*_sigX_*-*luxAB* reporter response in THY. C) P*_comS_*-*luxAB* reporter response in THY.

## DISCUSSION

Our study aimed to provide an assessment of basic parameters pertaining to the development of natural competence in *Streptococcus ferus*. We determined that *S. ferus* possesses a functional ComRS system but lacks a *S. mutans*-like BlpRH (formerly ComCDE) system. This mirrors what has been documented in some streptococci (14, 15, 18, 30) but the absence of BlpRH in *S. ferus* is somewhat surprising considering the close genetic relationship to *S. mutans* (21, 22). Addition of synthetic XIP derived from either *S. mutans* or *S. ferus comS* induced transcription of *comS* and *comX* and resulted in transformation. The compatibility of both XIP versions suggest that interspecies crosstalk and genetic transfer may occur if *S. ferus* and *S. mutans* encounter one other in the oral niche. We were motivated to investigate the genetic tractability of *S. ferus* due to our interest in the biosynthetic gene cluster (BGC) that produces tryglysin. We previously found that the *S. ferus* and *S. mutans* tryglysin BGCs are highly similar, and that the resulting products of these BGCs only differ by one amino acid (24). It therefore seems likely these genes were transferred horizontally between organisms as an outcome of interspecies crosstalk mediating genetic exchange (31, 32), and there is plausibility to the opportunity that *S. ferus* could acquire virulence traits from *S. mutans* that could lead to increased cariogenic potential.

As with other streptococcal species for which ComRS systems have been elucidated, *S. ferus* ComR is essential for inducing natural transformation (Fig. 2). This is not surprising, as this transcriptional regulator controls the production of SigX, the alternative sigma factor required for the development of competence. Also unsurprising is the presence of two *sigX* orthologs (Fig. 1) and we found that either is sufficient to allow for induction of competence, although disruption of either one alters transformation frequencies (Fig. 2, Fig. S2), also similar to that reported in other streptococci (7). The reason for two copies of *sigX* remains unclear although it may indicate the intense evolutionary pressure to conserve the ability to take up DNA.

As previously documented for *S. mutans* and *S. sobrinus*, *S. ferus* encodes an additional copy of *comR* in its genome (14, 30) In *S. mutans*, it is hypothesized that this secondary ComR regulates the production of downstream putative bacteriocins, but no published data has demonstrated this (14). S. *sobrinus* ComR1 and ComR2 are distinct in that they respond to an XIP that lacks aromatic residues but is highly enriched for hydrophobic amino acids, akin to signaling peptides for the closely related RRNPP family of QS-regulatory systems (reported C7 peptide LMCTIVR); and both have been reported to modulate competence (30, 33). In *S. ferus*, both *comR* and *comR2* are followed by predicted small ORFs that conserve the expected architecture for a *sigX*-regulated promoter (Fig. 1). Though *comS* is the prototypical XIP containing a double tryptophan motif, the predicted small ORFs downstream of *comR2* are unlike other characterized XIP or SHP peptides aside from their high enrichment for hydrophobic amino acids (Table 2).

In the preparation of this manuscript, a preprint examining the various aspects of *comR* orthologs in oral streptococci became available (Li *et al.,* 2022. *BioRxiV*). The authors report two *S. ferus comR* orthologs and *comS*, as we have. Differences between our report and Li *et al.* can be seen for ComR2, in that the *orf01* we describe is different from that described as XIP2b in the other study, although this region is conserved between the *S. ferus* strain we used (DSM20646) and the one in their study (8S1) (Fig. S4). We reason that among the 5 theoretical ORFs identified in the intergenic region, only two (*orf01* and *orf02*) would be transcribed from the promoters identified in Fig. 1C. Given the inherent difficulty to predict expressed sORFs (34), further investigation will be required to associate these coding sequences with physiological function. Interestingly, Li *et al*. found no impact of deletion of *comR2* or addition of the identified putative XIPs on *S. ferus* transformation (Li *et al.,* 2022. *BioRxiV*).

Of additional note is the impact that XIP appears to have on *S. ferus* physiology. At elevated concentrations of XIP, we observed diminished growth rates of *S. ferus* cultures that were not due to the vehicle solvent, nor seen from the nonsense peptide (revSHP) or XIP from *S. mutans* (Fig. 3). Interestingly, the slowed rate was abrogated when both *sigX* genes were deleted, indicating that induction of competence is required for this phenotype (Fig. S3). Competence induction is likely to place an acute physiological stress on the cell and the cell cycle may be interrupted in *S. ferus*, as has been described for *S. pneumoniae* and *B. subtilis*, although distinct mechanisms have been elucidated in these systems (35, 36). Induction of competence may also lead to the production of bacteriocins or fratricidal products, like that observed in *S. mutans* via ComCDE (alternatively known as BlpCHR or MutCDE) or for *S. pneumoniae* with its fratricidal competence induced behavior (2, 3, 11, 19, 20). It is currently unknown if any other bacteriocins or fratricidal compounds aside from tryglysin A are produced by *S. ferus* and further examination will be required to examine these hypotheses. Nevertheless, our work provides a foundation for further genetic manipulation of *S. ferus* provides a framework for a simple model of competence and potential crosstalk with other species.

## MATERIALS AND METHODS

### Bacterial strains, plasmids, and growth conditions

Bacterial strains and plasmids used in this study are listed in Table S1. *S. ferus* strains were derived from DSM20646 (37) were grown on Todd-Hewitt plates (TH, BD Biosciences) with 1.4% Bacto-Agar (BD Biosciences) and 0.2% yeast extract (VWR), in Todd-Hewitt broth with 0.2% yeast (THY), or in a chemically-defined medium (CDM) containing 1% glucose at 37°C with 5% CO_2_. The components and recipe for CDM used were as described previously (38). *E. coli* strains were grown on Luria-Bertani (BD Biosciences) plates with 1.4% Bacto-Agar at 37°C or in Luria-Bertani (LB) broth with shaking at 175 rpm. When required spectinomycin (50-100 µg/mL), erythromycin (0.5 µg/mL for *S. ferus*, 500 µg/mL for *E. coli*), or kanamycin (50-100 µg/mL) were added to *S. ferus* or *E. coli* culture media.

### Identification of ComR, ComS, SigX, and Rgg orthologs in *S. ferus*, search for ComCDE orthologs and subsequent sequence analysis

To determine if proteins like *S. mutans* ComR, ComR2, SigX, PdrA (another Rgg), and ComCDE (aka BlpRH) were present in *S. ferus,* the amino acid sequences of each *S. mutans* UA159 protein were input into the online NCBI protein BLAST to identify potential orthologs (25). The ortholog for *S. mutans* PdrA (a Rgg that regulates production of a RaS-RiPP biosynthetic operon) has been previously identified in *S. ferus* as A3GY_ RS0100490 (24, 37). These searches were performed using the following parameters: Database, Non-redundant protein sequences (nr); Organism, *Streptococcus ferus* (taxid:1345); Algorithm, blastp (protein-protein BLAST); all other parameters set to default. The following amino acid sequences and their respective genetic loci from *S. mutans* UA159 were used: ComR, WP_002263400.1, *SMU_61*; ComR2, WP_002262469.1, *SMU_381c*; SigX, WP_002263585.1, *SMU_1997*; PdrA, WP_002352338.1, *SMU_1509*; BlpC (formerly ComC), WP_002267610.1, *SMU_1915*; BlpH (formerly ComD), WP_002262113.1, *SMU_1916*; BlpR (formerly ComE), WP_002262114.1, *SMU_1917*. Identified hits to *S. mutans* ComR, ComR2, PdrA, and SigX in *S. ferus* DSM20646 (Assembly: GCA_000372425.1) were then input alongside the original *S. mutans* sequences into the Clustal Omega webserver (https://www.ebi.ac.uk/Tools/msa/clustalo/) using default parameters (39) to obtain multiple amino acid sequence alignment and percent identity for each comparison. No orthologs to *S. mutans* BlpC, BlpH, or BlpR were identified via protein BLAST. Upon further inspection, we determined that the assembly for DSM20646 was highly similar if not identical to the NCTC2278 assembly (Assembly: GCA_900475025.1) listed in NCBI genome, as although these have been annotated as separate strains in NCBI genome are annotated as the same strain in the National Collection of Type Cultures from the U.K. Health Security Agency (https://www.culturecollections.org.uk).

To identify the putative ComS orthologs in *S. ferus*, the genetic architecture surrounding the identified ComR ortholog (A3GY_RS0108865) was examined for short open reading frames using SnapGene 6.1.2 (SnapGene Software, www.snapgene.com) setting the translation options to require a minimum ORF length of 10 amino acids and a ATG start codon. Identified short open reading frames 3’ after *comR* were prioritized, as *comS* directly follows *comR* in *S. mutans* (14). This identified putative open reading frames after *comR* were then examined for promoter architecture by input of 200 bp of nucleotides from the 5’ sequence directly upstream of the identified ORFs into the online BPROM server using default parameters (http://www.softberry.com/berry.phtml?topic=bprom&group=programs&subgroup=gfindb) (40). Predicted promoter architecture was then manually compared to the known *comS* promoter sequence from *S. mutans* UA159. The small open reading frame most closely matching the *S. mutans comS* promoter architecture and a putative double tryptophan motif was identified as a short open reading frame of 13 amino acids 76 bp from the stop codon of A3GY_RS0108865 and was designated as the putative *S. ferus comS*. We examined the architecture around another identified BLAST hit to *S. mutans* ComR, A3GY_RS0106270 in a similar manner and were unable to identify a short open reading frame with a double tryptophan motif near this gene but did identify a similar promoter architecture to *comS* in front of two sORFs that appeared to encode for two separate peptides, as shown in Fig. 1 and Table 2. In addition to these two sORFs, three other putative sORFs were found in this region (Fig. S4A).

### Construction of linear DNA amplicons for *S. ferus* transformation

To obtain linear DNA amplicons for *S. ferus* transformation, *S. ferus* genomic DNA or *E. coli* plasmid DNA were amplified by PCR with Phusion High-Fidelity DNA Polymerase (New England Biolabs, NEB) using primers as indicated in Table S1. Fragments were then purified with the DNA Clean & Concentrator kit (Zymo Research) and assembled via Gibson assembly using NEBuilder HiFi DNA Assembly MasterMix (NEB) according to the manufacturer’s instructions. Resulting Gibson assemblies were then amplified again by PCR using flanking primers to confirm correct size of assembly before construction of strains as detailed in *Transformation of S. ferus and construction of S. ferus strains*.

For construction of *comS*(GGG)-*spec* (also listed as *comS*(ATG→GGG)-*spec* in Table S1: DNA fragments mutated for the start codon of *comS*(ATG→GGG) were obtained from *S. ferus* wild-type genomic DNA by PCR with primers BR435/BR523 and BR522/BR436. Fragments were purified as detailed above and then fused by PCR using 1 µL each as a template with primers BR435/BR436. This fragment was designated as the *comS*(GGG) fusion amplicon, and 5’ and 3’ fragments for final construction of *comS*(GGG)-*spec* were amplified from this amplicon with primers BR435/BR524 and BR527/BR436. The *spec* cassette was obtained from genomic DNA extracted from strain BRSF04 using primer set BR525/BR526, and all fragments were purified. The resulting fragments were ligated together via Gibson assembly using NEBuilder HiFi DNA Assembly MasterMix according to the manufacturer’s instructions and verified for correct assembly by PCR before being used for transformation.

### Construction of plasmids for *S. ferus* transformation

All plasmids were constructed via Gibson assembly of DNA fragments using NEBuilder HiFi DNA Assembly MasterMix according to the manufacturer’s instructions. All DNA fragments were amplified by PCR and purified as detailed in *Construction of linear DNA amplicons for S. ferus transformation.* Resulting plasmid assemblies were electroporated into *E. coli* DH5α using a Bio-Rad Gene Pulser II Electroporation system (Bio-Rad) with the following parameters: 2.0 V, 200 Ω, 250 µF. Constructed plasmids were extracted using a GeneJET Plasmid MiniPrep Kit (Thermo-Fisher) according to the manufacturer’s instructions and confirmed by restriction digest, PCR, and Sanger sequencing. All plasmids were propagated in DH5α cells for further use. Primers and additional information are listed in Table S1.

### Synthesis of *S. ferus* XIP, *S. mutans* XIP, and *S. mutans* reverse SHP peptides

Synthetic peptides were purchased from ABClonal (Woburn, MA) or NeoScientific (Cambridge, MA). Purities of peptide preparations used in assays were greater than 70%. All peptides were reconstituted as 10 mM stocks in DMSO and stored in aliquots at −20°C. For *S. ferus* XIP and *S. mutans* XIP, subsequent dilutions for working stocks (1 mM, 100 µM, 10 µM) were made in DMSO and stored at −20°C.

### Transformation of *S. ferus* and construction of *S. ferus* strains

For transformation, *S. ferus* strains were grown overnight from frozen glycerol stocks in consecutive THY dilutions at 37°C with 5% CO_2_. The next morning, overnight strains were spun down at 4000*xg* for 10 min, and subsequently resuspended in 1 mL of prewarmed CDM. Alternatively, starter cultures of strains stored at −70°C in CDM plus 20% glycerol were used. Overnight strains or starter cultures were spun at 11,000*xg* for 5 min, and resuspended again in 1 mL of CDM. Resulting resuspended strains were then diluted to OD_600_ ∼0.05 in prewarmed CDM, in which cultures were then split into 500 µL aliquots for separate transformation conditions individually or in triplicate. Cultures were then incubated for 1 hour at 37°C with 5% CO_2_, at which time 1 µM of *S. ferus* or *S. mutans* XIP and 50-100 ng of DNA were added to required tubes. A negative control was performed for all strains/conditions as well, in which either no DNA or no DNA/XIP was added. Cultures were placed back at 37°C with 5% CO_2_ for two hours, at which time cultures were serially diluted and plated for CFU on THY and THY with required antibiotic for selection. Plates were then incubated at 37°C with 5% CO_2_ for 1-3 days before colonies were enumerated. It was noted that colonies were most visible and easiest to count at 2 days post incubation but that absolute numbers did not change if colonies were counted from 1-3 days post inoculation. Resulting CFU/ml for each condition and replicate were then used to calculate transformation efficiency and frequency as follows. Transformation efficiency (TE) was calculated as (TE = average number of transformant colonies/µg DNA/dilution of transformants). Transformation frequency (TF) was calculated as (TF = (CFU/mL of transformants / total CFU/mL) x 100). To examine significant differences between transformation conditions, final TE and TF were input into Graph Pad Prism 9.5.0. (GraphPad Software, LLC) and examined for statistical significance via an Unpaired T test (if only 2 conditions were examined) or a One-way ANOVA with Tukey’s Multiple Comparisons Post-test (for more than 2 conditions).

To examine the effect of DNA concentration, XIP concentration, and time on transformation efficiency, transformations were performed as above, with the following modifications. For examination of the effect of DNA concentration, wild-type *S. ferus* samples in triplicate were transformed with increasing amounts of linear Δ*comR*::*spec* amplicon (0 ng, 5 ng, 10 ng, 50 ng, 100 ng, and 200 ng total DNA) with XIP held constant at 1 µM. A negative control transformation was also performed with no XIP or DNA. To examine effect of XIP concentration, wild-type *S. ferus* samples in triplicate were transformed with increasing amounts of *S. ferus* XIP (0 nM, 10 nM, 100 nM, 1 µM, 10 µM), with linear Δ*comR*::*spec* DNA amount held constant at 100 ng. A negative control transformation was also performed with no XIP or DNA. To examine the effect of time on transformation, wild-type *S. ferus* samples in triplicate were transformed with 50 ng of linear Δ*comR*::*spec* DNA and 1 µM XIP and plated at various times post XIP/DNA addition (0 min, 5 min, 15 min, 30 min, 45 min, 1 hr, 2 hr, and 3 hr). A control transformation with no DNA or XIP was also performed alongside this experiment and plated at 3 hours post experiment start.

If transformations were performed to obtain strains for later use, single colonies were isolated and subsequently stored overnight in THY plus required antibiotic. The next morning, single isolates were stored with 20% glycerol at −70°C. A portion of the resulting culture was then spun down at 4000*xg* for 10 min, and genomic DNA was extracted using a MasterPure Gram Positive DNA Purification Kit (Epicentre, MGP04200) according to the manufacturer’s instructions or, if containing plasmids requiring verification, according to the published protocol by Freitas *et al.*, 2020 for *Extraction of Low/Medium MW Plasmids by an Alkaline Lysis-Based Method* (41) or alternatively by digestion of *S. ferus* cells and extraction with the GeneJET Plasmid MiniPrep Kit as follows. To purify plasmid from *S. ferus* using a GeneJET Plasmid MiniPrep Kit, overnight cultures of *S. ferus* strains in THY were spun down at 4000*xg* for 10 min. Strains were resuspended in digestion mix (150 µL Tris-EDTA, pH 8.0, 1 µL of Ready-Lyse Lysozyme Solution from a MasterPure Gram Positive DNA Purification Kit (Epicentre), 45 units mutanolysin) and incubated at 37°C for 1.5 hrs. Cells were then processed as according to manufacturer’s instructions for the GeneJET Plasmid MiniPrep Kit. Resulting genomic DNA or plasmid DNA was used to confirm strains via PCR and Sanger sequencing.

### Test ability of stationary phase supernatants to induce transformation

Overnight cultures of wild-type *S. ferus* were grown in 6 mL of CDM or THY at 37°C with 5% CO_2_. The next morning, the cultures were centrifuged at 4000*xg* for 10 min. The culture grown in CDM was sterile filtered with a 25 mm 0.22 µM PES filter (VWR) and transferred to a new test tube. The culture grown in THY was resuspended in 1 mL of prewarmed CDM. The culture was spun again at 13,000*xg* for 5 min, and resuspended again in 1 mL of CDM. After this, the resuspended culture was diluted to OD_600_ ∼0.05 in the 6 mL of filtered supernatant or 6 mL of fresh prewarmed CDM. Resulting cultures were aliquoted at 500 µL in Eppendorf tubes and incubated for 1 hour at 37°C with 5% CO_2_ before exposure to the following conditions. For fresh prewarmed CDM cultures: 100 ng of DNA and 1 µM *S. ferus* XIP, No DNA and 1 µM *S. ferus* XIP, No DNA and No XIP. For filtered supernatant: 100 ng of DNA and 1 µM *S. ferus* XIP, No DNA and No XIP. Cultures were incubated for 2 hours after exposure and plated for on THY and THY with required antibiotic for selection. Plates were then incubated at 37°C with 5% CO_2_ for 2-3 days before colonies were enumerated. TE, TF, and statistical significance were calculated, graphed, and observed as detailed in *Transformation of S. ferus and construction of S. ferus strains*.

### Growth curves of strains

Overnight cultures grown in THY at 37°C with 5% CO_2_ or starter cultures of strains stored at −70°C in CDM plus 20% glycerol were washed in prewarmed CDM. For overnight cultures, strains were spun down at spun down at 4000*xg* for 10 min, resuspended in 1 mL of prewarmed CDM, centrifuged again at 11,000*xg* for 5 min, and resuspended again in 1 mL of CDM. For starter cultures already in CDM, strains were centrifuged at 11,000*xg* for 5 min and resuspended in 1 mL of CDM. Strains were then resuspended in 6 mL prewarmed CDM to OD_600_ ∼0.05 and grown at 37°C with 5% CO_2_, with OD_600_ observed every 45 min to 1 hour with a GENESYS 30 Vis spectrophotometer (Thermo-Fisher). At 1 hour post inoculation (approximately OD_600_ ∼0.05-0.1), 10 nM, 100 nM, or 1 µM of *S. ferus* XIP, *S. mutans* XIP, *S. mutans* reverse SHP, or an equivalent volume of DMSO was added to cultures. OD_600_ was continued to be observed by spectrophotometer every 45 min to 1 hour after this addition. Resulting data from growth curves was plotted and doubling times were calculated by taking absorbance values from OD_600_ ∼0.1 – 0.9 and performing a Nonlinear regression analysis with Exponential growth equation fitting with Graph Pad Prism 9.5.0. Statistical significance between doubling times were determined by One-way ANOVA with Dunnett’s or Šidák’s Multiple Comparisons Post-test with Graph Pad Prism 9.5.0.

### Luciferase assays

Overnight cultures were grown and washed with prewarmed CDM as described in *Growth curves of strains.* At 1 hour post inoculation (approximately OD_600_ ∼0.05-0.1), approximately 10 nM, 100 nM, or 1 µM *S. ferus* XIP, *S. mutans* XIP, or *S. mutans* reverse SHP were added to cultures. Every 30 min to 1 hour, OD_600_ measurements were taken with a GENESYS 30 Vis spectrophotometer and luciferase measurements were conducted by transferring 100 µL aliquots from each condition/strain to an opaque 96-well plate. Samples were exposed to decyl aldehyde (Sigma-Aldrich) fumes for one minute and counts per second (CPS) were measured using a Veritas microplate luminometer (Turner Biosystems). Relative light units (RLU) were calculated by normalizing CPS to OD_600_. Data from experiments were plotted and analyzed using Graph Pad Prism 9.5.0.

## ACKNOWLEDGEMENTS

The authors would like to thank the members of the Federle lab for helpful discussions and proofreading of the manuscript.

## FUNDING AND ADDITIONAL INFORMATION

This study was supported by the National Institutes of Health (R01AI091779 to M.J.F. and 5K12 GM139186/NIGMS/NIH to B.E.R.).

## REFERENCES

1. Dubnau D, Blokesch M. 2019. Mechanisms of DNA Uptake by Naturally Competent Bacteria. Annu Rev Genet 53:217–237.

2. Straume D, Stamsås GA, Håvarstein LS. 2015. Natural transformation and genome evolution in *Streptococcus pneumoniae*. Infection, Genetics and Evolution 33:371–380.

3. Salvadori G, Junges R, Morrison DA, Petersen FC. 2019. Competence in streptococcus pneumoniae and close commensal relatives: Mechanisms and implications. Front Cell Infect Microbiol https://doi.org/10.3389/fcimb.2019.00094.

4. Winter M, Buckling A, Harms K, Johnsen PJ, Vos M. 2021. Antimicrobial resistance acquisition via natural transformation: context is everything. Curr Opin Microbiol 64:133– 138.

5. Håvarstein LS, Coomaraswamy G, Morrison DA. 1995. An unmodified heptadecapeptide pheromone induces competence for genetic transformation in *Streptococcus pneumoniae*. Proceedings of the National Academy of Sciences 92:11140–11144.

6. Håvarstein LS, Gaustad P, Nes IF, Morrison DA. 1996. Identification of the streptococcal competence-pheromone receptor. Mol Microbiol 21:863–869.

7. Lee MS, Morrison DA. 1999. Identification of a New Regulator in *Streptococcus pneumoniae* Linking Quorum Sensing to Competence for Genetic Transformation. J Bacteriol 181:5004–5016.

8. Morrison DA, Lee MS. 2000. Regulation of competence for genetic transformation in *Streptococcus pneumoniae*: a link between quorum sensing and DNA processing genes. Res Microbiol 151:445–451.

9. Peterson S, Cline RT, Tettelin H, Sharov V, Morrison DA. 2000. Gene Expression Analysis of the *Streptococcus pneumoniae* Competence Regulons by Use of DNA Microarrays. J Bacteriol 182:6192–6202.

10. Peterson SN, Sung CK, Cline R, Desai B v., Snesrud EC, Luo P, Walling J, Li H, Mintz M, Tsegaye G, Burr PC, Do Y, Ahn S, Gilbert J, Fleischmann RD, Morrison DA. 2004. Identification of competence pheromone responsive genes in *Streptococcus pneumoniae* by use of DNA microarrays. Mol Microbiol 51:1051–1070.

11. Shanker E, Federle MJ. 2017. Quorum sensing regulation of competence and bacteriocins in *Streptococcus pneumoniae* and *mutans*. Genes (Basel) https://doi.org/10.3390/genes8010015.

12. Li Y-H, Lau PCY, Lee JH, Ellen RP, Cvitkovitch DG. 2001. Natural Genetic Transformation of *Streptococcus mutans* Growing in Biofilms. J Bacteriol 183:897–908.

13. Dhaked HPS, Biswas I. 2022. Distribution of two-component signal transduction systems BlpRH and ComDE across streptococcal species. Front Microbiol 13:960994.

14. Mashburn-Warren L, Morrison DA, Federle MJ. 2010. A novel double-tryptophan peptide pheromone is conserved in mutans and pyogenic Streptococci and Controls Competence in *Streptococcus mutans* via an Rgg regulator. Mol Microbiol 78:589–606.

15. Fontaine L, Boutry C, de Frahan MH, Delplace B, Fremaux C, Horvath P, Boyaval P, Hols P. 2010. A Novel Pheromone Quorum-Sensing System Controls the Development of Natural Competence in *Streptococcus thermophilus* and *Streptococcus salivarius*. J Bacteriol 192:1444–1454.

16. Gardan R, Besset C, Guillot A, Gitton C, Monnet V. 2009. The Oligopeptide Transport System Is Essential for the Development of Natural Competence in *Streptococcus thermophilus* Strain LMD-9. J Bacteriol 191:4647–4655.

17. Talagas A, Fontaine L, Ledesma-Garca L, Mignolet J, Li de la Sierra-Gallay I, Lazar N, Aumont-Nicaise M, Federle MJ, Prehna G, Hols P, Nessler S. 2016. Structural Insights into Streptococcal Competence Regulation by the Cell-to-Cell Communication System ComRS. PLoS Pathog 12:e1005980.

18. Shanker E, Morrison DA, Talagas A, Nessler S, Federle MJ, Prehna G. 2016. Pheromone Recognition and Selectivity by ComR Proteins among Streptococcus Species. PLoS Pathog 12:e1005979.

19. Lemos JA, Palmer SR, Zeng L, Wen ZT, Kajfasz JK, Freires IA, Abranches J, Brady LJ. 2019. The Biology of *Streptococcus mutans*. Microbiol Spectr 7:10.1128/microbiolspec.GPP3-0051–2018.

20. Kaspar JR, Walker AR. 2019. Expanding the Vocabulary of Peptide Signals in *Streptococcus mutans*. Front Cell Infect Microbiol 9:194.

21. Freedman ML, Coykendall AL, O’Neill EM. 1982. Physiology of “mutans-like” *Streptococcus ferus* from wild rats. Infect Immun 35:476–82.

22. Baele M, Devriese LA, Vancanneyt M, Vaneechoutte M, Snauwaert C, Swings J, Haesebrouck F. 2003. Emended description of *Streptococcus ferus* isolated from pigs and rats. Int J Syst Evol Microbiol 53:143–146.

23. Igarashi M. 2008. Identification of *Streptococcus ferus* and the Distribution of Three Kinds of Mutants Streptococci from Pig Tooth Surface. International Journal of Oral-Medical Sciences 7:7–11.

24. Rued BE, Covington BC, Bushin LB, Szewczyk G, Laczkovich I, Seyedsayamdost MR, Federle MJ. 2021. Quorum Sensing in *Streptococcus mutans* Regulates Production of Tryglysin, a Novel RaS-RiPP Antimicrobial Compound. mBio 12: e02688–20.

25. Altschul SF, Gish W, Miller W, Myers EW, Lipman DJ. 1990. Basic local alignment search tool. J Mol Biol 215:403–410.

26. Opdyke JA, Scott JR, Moran, Charles P. 2003. Expression of the Secondary Sigma Factor σ ^X^ in *Streptococcus pyogenes* Is Restricted at Two Levels. J Bacteriol 185:4291– 4297.

27. Mashburn-Warren L, Morrison DA, Federle MJ. 2012. The Cryptic Competence Pathway in *Streptococcus pyogenes* Is Controlled by a Peptide Pheromone. J Bacteriol 194:4589– 4600.

28. Contente S, Dubnau D. 1979. Characterization of plasmid transformation in *Bacillus subtilis*: Kinetic properties and the effect of DNA conformation. Mol Gen Genet 167:251– 258.

29. Knapp S, Brodal C, Peterson J, Qi F, Kreth J, Merritt J. 2017. Natural Competence Is Common among Clinical Isolates of *Veillonella parvula* and Is Useful for Genetic Manipulation of This Key Member of the Oral Microbiome. Front Cell Infect Microbiol 7:139.

30. Li JW, Wyllie RM, Jensen PA. 2021. A Novel Competence Pathway in the Oral Pathogen *Streptococcus sobrinus*. J Dent Res 100:542–548.

31. Cook LC, LaSarre B, Federle MJ. 2013. Interspecies communication among commensal and pathogenic streptococci. mBio 4:e00382–13.

32. Milly TA, Tal-Gan Y. 2020. Biological evaluation of native streptococcal competence stimulating peptides reveals potential crosstalk between *Streptococcus mitis* and *Streptococcus pneumoniae* and a new scaffold for the development of *S. pneumoniae* quorum sensing modulators. RSC Chem Biol 1:60–67.

33. Cook LC, Federle MJ. 2014. Peptide pheromone signaling in *Streptococcus* and *Enterococcus*. FEMS Microbiol Rev 38:473–492.

34. Laczkovich I, Mangano K, Shao X, Hockenberry AJ, Gao Y, Mankin A, Vázquez-Laslop N, Federle MJ. 2022. Discovery of Unannotated Small Open Reading Frames in *Streptococcus pneumoniae* D39 Involved in Quorum Sensing and Virulence Using Ribosome Profiling. mBio 13: e0124722.

35. Bergé MJ, Mercy C, Mortier-Barrière I, VanNieuwenhze MS, Brun Y v., Grangeasse C, Polard P, Campo N. 2017. A programmed cell division delay preserves genome integrity during natural genetic transformation in *Streptococcus pneumoniae*. Nat Commun 8:1621.

36. Hahn J, Tanner AW, Carabetta VJ, Cristea IM, Dubnau D. 2015. ComGA-RelA interaction and persistence in the *Bacillus subtilis* K-state. Mol Microbiol 97:454–471.

37. Bushin LB, Clark KA, Pelczer I, Seyedsayamdost MR. 2018. Charting an Unexplored Streptococcal Biosynthetic Landscape Reveals a Unique Peptide Cyclization Motif. J Am Chem Soc 140:17674–17684.

38. Chang JC, LaSarre B, Jimenez JC, Aggarwal C, Federle MJ. 2011. Two group a streptococcal peptide pheromones act through opposing rgg regulators to control biofilm development. PLoS Pathog 7:e1002190.

39. Sievers F, Wilm A, Dineen D, Gibson TJ, Karplus K, Li W, Lopez R, McWilliam H, Remmert M, Söding J, Thompson JD, Higgins DG. 2011. Fast, scalable generation of high-quality protein multiple sequence alignments using Clustal Omega. Mol Syst Biol 7:539.

40. Solovyev V, Salamov A. 2011. Automatic annotation of microbial genomes and metagenomic sequences, p. 61–78. In Metagenomics and its Applications in Agriculture, Biomedicine and Environmental Studies.

41. Freitas AR, Novais C, Peixe L, Coque TM. 2020. Isolation and Visualization of Plasmids from Gram-Positive Bacteria of Interest in Public Health, p. 21–38. In Methods Mol Biol.

